# A family-based phasing algorithm for sequence data

**DOI:** 10.1101/504480

**Authors:** Mara Battagin, Serap Gonen, Roger Ros-Freixedes, Andrew Whalen, Gregor Gorjanc, John M Hickey

## Abstract

This paper describes a family-based phasing algorithm, for variable-coverage sequence data, that first minimises phasing errors and then maximises the proportion of alleles phased. This algorithm is one of the essential tools that underpin an overall strategy for generating highly accurate sequence data on whole populations at low cost.

The algorithm is called AlphaFamSeq. It uses sequence data on the focal individual and at least two generations of ancestors to phase alleles. In the first step, AlphaFamSeq calculates allele probabilities using iterative peeling. In subsequent steps, the alleles are phased using heuristics deriving information from the sequence data of parents, grandparents and progenies and, if available, from other families in the pedigree. AlphaFamSeq was tested on a range of simulated data sets.

AlphaFamSeq gives low phasing error rates and, if there is sufficient sequence information and haplotype sharing amongst individuals, it can give a high yield of correctly phased alleles.

The allele threshold had a large effect and window size had a small effect on performance. When all individuals in a single family were sequenced at different coverages the highest correctly phased alleles reached 90% of the possible maximum (98.9%) at ~1/6 of the maximum aggregate coverage. Adding sequence information from other related individuals increased the percentage of correctly phased alleles. Imputation performance was high across all allele frequencies (average correlation by marker of 0.94), except for a slight decrease at very low frequencies (≤0.01 MAF).

Within an overall strategy for generating highly accurate sequence data on whole populations at low cost the role of AlphaFamSeq is to provide very accurately phased haplotypes on focal individuals, who are individuals whose haplotypes are very common in the population.

## Background

This paper describes a family-based phasing algorithm for variable-coverage sequence data that first minimises phasing errors and then maximises the proportion of alleles phased. This design enables accurate imputation of sequence data in livestock populations. Superficially the work of Robin Thompson has tenuous links to phasing of sequence data. He was the first to recognise that the statistical modelling of breeding values could be partitioned into parent average and Mendelian sampling terms. Phasing, imputation and sequencing strategies exploit the same genetic principles. This paper describes one such method.

In a livestock population, sequence data has a number of potential advantages compared to classical marker genotype data including increased power to discover causative variants and more accurate genomic predictions. Sequence data has enabled the discovery of causative variants for qualitative traits (e.g. for embryonic lethality in the 1,000 Bulls Project [1]). However, sequence data has only shown modest benefits for quantitative traits, and a small increases in the accuracy of genomic prediction [2,3].

There are two likely reasons for the lack of large benefit for quantitative traits. First, data from millions of individuals may be required to capture the full potential of sequence data in livestock [4]. With fewer individuals, there will not be enough recombination events to estimate the effects of the large numbers of causative variants that underlie a quantitative trait. Second, the imputation methods used to generate large datasets may not be accurate enough. There may not be enough information in the generated sequence data to enable accurate phasing. Existing phasing algorithms are sub-optimal because the animals that provide the sequence data are insufficiently related to the animals that have sequence data imputed.

We believe that these issues can be addressed if effective sequencing strategies and accurate phasing/imputation algorithms are developed jointly, which requires strategies for the optimal distribution of sequence resources across a population and accurate phasing/imputation algorithms. Separate strategies and algorithms are needed for low coverage sequence data and for variable coverage sequence data. We have recently developed AlphaSeqOpt, which implements the required strategies for the optimal distribution of sequence resources [5]. Now we need accurate phasing algorithms, specifically designed to meet the requirements of AlphaSeqOpt.

All imputation-based sequencing strategies involve: (i) collecting sequence data on a relatively small number of individuals; (ii) accurately phasing these individuals to resolve their haplotypes; and (iii) imputing the phased haplotypes into a large number of unsequenced individuals by inferring the combination of haplotypes that each unsequenced individual carries. In this context, the accurate phasing of the haplotypes is essential: phasing errors lead to incorrect imputation. In populations with hundreds of thousands of individuals this leads to large numbers of errors because haplotypes are carried by thousands of individuals.

Many phasing algorithms or analogous imputation algorithms have been developed (e.g., AlphaImpute [6], Findhap [7], Fimpute [8]) or used (e.g., MaCH [9], Beagle [10,11], Impute2 [12]) in livestock populations. These algorithms were originally developed for SNP genotype data, but some can also be used for sequence data. However, we believe that these algorithms are suboptimal, because they do not prioritise accurate phasing. Instead they seek to phase and impute alleles for all individuals at all genome positions. This approach generates good phasing for most of the markers, but fails in phasing low frequency alleles, which are common in sequence data sets. The error rates are especially high in algorithms that preferentially determine phase and impute alleles based on haplotypes that have high frequency (i.e., MaCH [9], Beagle [10,11], Impute2 [12], Findhap [7], Fimpute [8]).

The objective of this research was to develop a family-based phasing algorithm for variable-coverage sequence data that first minimises phasing errors and then maximises the proportion of alleles phased. This algorithm was designed to work with the sequencing strategy proposed by Gonen et al [5], which distributes a fixed amount of sequencing resources across a population by identifying focal individuals, determining the genomic footprint of these focal individuals and determining the optimal distribution of sequencing resources across these focal individuals and their ancestors (parents and grandparents) according to the genomic footprint of the focal individuals on the population.

The resulting algorithm performs well. It can work within a given family separately, can work with multiple families simultaneously by utilising pedigree information that connects them and can handle variable coverage sequencing data. The algorithm was implemented in a new software package called AlphaFamSeq. It gives low phasing error rates and, if there is sufficient sequence information and haplotype sharing amongst individuals, it can give a high yield of correctly phased alleles.

## Methods

We developed and tested a family-based phasing algorithm for variable-coverage sequence data that first minimises phasing errors and then maximises the proportion of alleles phased. The algorithm is called AlphaFamSeq; it uses sequence data on the focal individual and a pedigree that includes at least the parents and grandparents of the focal individual and requires that some of the individuals in this pedigree have sequence data. AlphaFamSeq processes individuals from the oldest to the youngest and iterates until convergence. In the first step, AlphaFamSeq calculates allele probabilities using iterative peeling [13]. In subsequent steps, the genotypes are phased using heuristics that derive information from the genomic data of grandparents, parents, and progeny and, if available, from other families in the pedigree.

In the next sections we give a detailed description of the data used by AlphaFamSeq, the algorithm and its parameters, the simulation of the data used to test the algorithm, and the metrics used in the tests.

### Data used by AlphaFamSeq

AlphaFamSeq uses pedigree information and genomic data, which can be sequence data or genotype data. Sequencing data can cover the whole genome or parts of it. The latter is common with the reduced representation sequencing approaches such as genotyping-by-sequencing [14]. The pedigree information must include the parents and grandparents of at least one focal individual. There is no upper limit on pedigree complexity. Within the pedigree individuals can be sequenced to any coverage or unsequenced. The sequence data are presented to the algorithm in the form of the number of reads for the reference and the alternative alleles at each bi-allelic variant.

### The AlphaFamSeq algorithm

AlphaFamSeq iterates across five main steps until convergence. These steps are summarised in Figure 1.

**Figure1.**
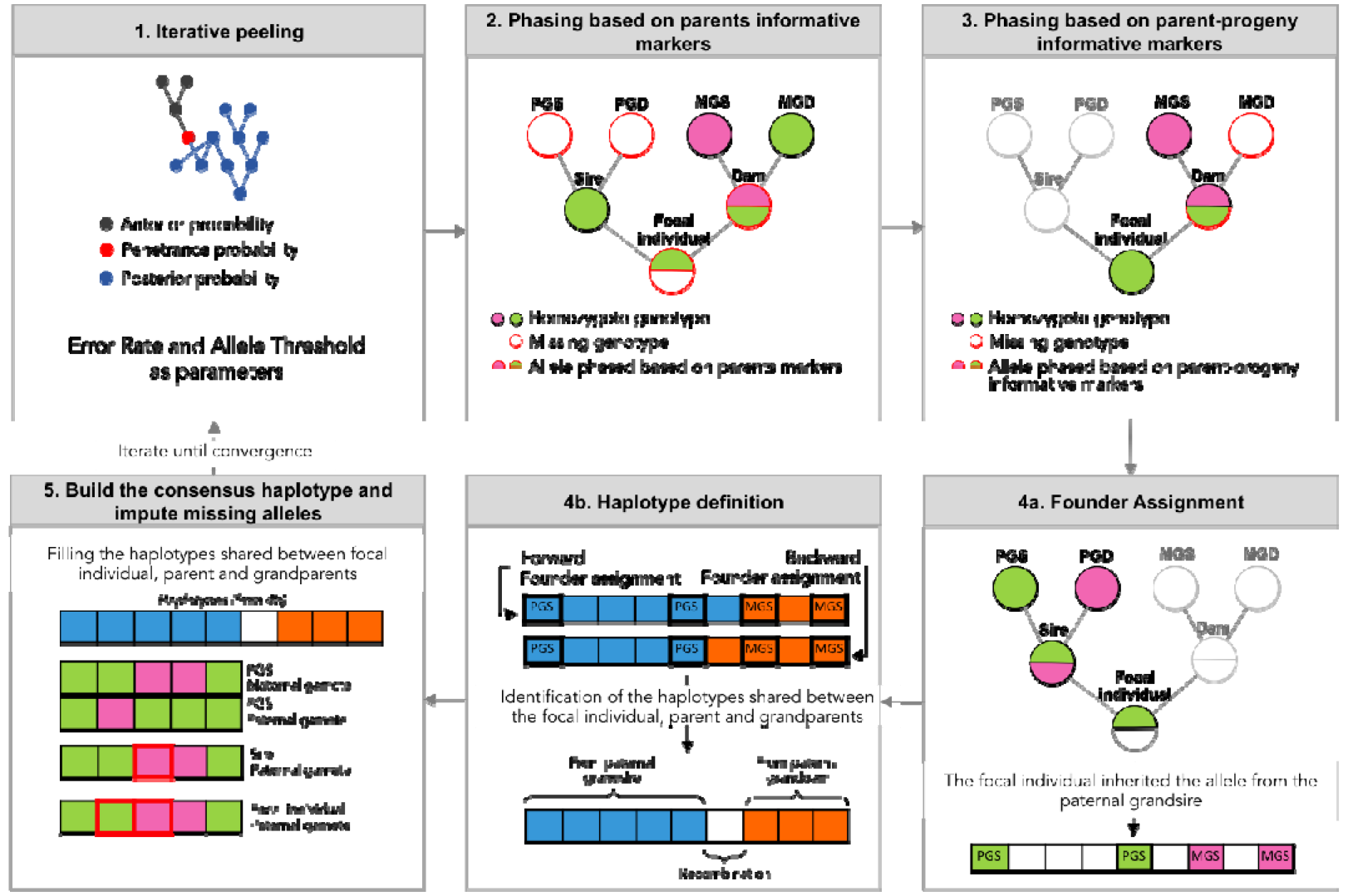
AlphaFamSeq steps. PGS = Paternal Grandsire, PGD= Paternal Granddam, MGS = Maternal Grandsire, MGD= Maternal Granddam

#### 1. Iterative peeling

The first step calculates allele probabilities for each variant of each individual in the pedigree, using all the pedigree information and the genomic data available. This step is based on a modified version of the iterative peeling method described by Kerr and Kinghorn [13]. The penetrance function is modified so that it accepts both observed genotypes or observed number of reads for the reference and the alternative alleles. The penetrance function is based on the binomial distribution with a given number of trials (number of sequence reads) and user-defined error rate [9]. This method accumulates information from parents, the individual itself, and progeny (accounting for mates’ information), which enables propagation of genotype or sequence information across large and complex pedigrees typical of livestock populations [13]. The peeling step is a single locus phasing where the outputs are allele probabilities. If an allele probability, is above a user-defined threshold the allele is called. In AlphaFamSeq the peeling is performed once, in the first iteration, and its output is used differently at each subsequent iteration by dynamically relaxing the user-defined allele thresholds. This allows the algorithm to utilise only the very informative variants in the early iterations to ensure that errors are not generated and subsequently propagated but to relax this stringency in later iterations to enable the yield to increase.

#### 2. Phasing based on parents informative markers

In the second step a Mendelian inheritance rule is used to phase alleles of an individual if one or both of its parents are deemed homozygous either from sequencing reads or from information derived from a previous iteration or a previous step. The rule is: if a parent is homozygous at a variant, then the phase of the progeny for the gamete inherited from that parent can be assigned.

#### 3. Phasing based on parent-progeny informative markers

In the third step, a Mendelian inheritance rule is used to phase alleles of an individual on the basis of marker information gathered from parent to progeny inheritance. The rule is: if an individual has one of its two gametes phased and the evidence from the progeny indicates that it inherited the other allele, then on the basis of progeny information the allele can be phased for the relevant gamete of the parent.

#### 4. Founder Assignment and haplotype definition

In the fourth step, rules are used to identify trios of individuals (a focal individual, its relevant parent and grandparent) that share a haplotype. The grandparents are considered the founders (within the context of this three generation sub-pedigree) and the aim is to assign the correct founder to each allele in the focal individual. In this step AlphaFamSeq allows the user to divide the chromosome into different windows. Windows are used in AlphaFamSeq for two reasons:

i. to speed up the steps 4 and 5 of AlphaFamSeq by working on multiple windows in parallel; and
ii. to limit the search space when individuals are sequenced at low-coverage and there are not enough informative variants to detect recombinations.

At each heterozygous locus of the parent, if the allele that the focal individual inherited from that parent is known, the grandparent from which the focal individual inherited the allele is determined. Once this is complete for each locus, each gamete of the focal individual is examined from the beginning to the end and from the end to the beginning of a user defined window. If the same founder is identified across the entire window by the examinations in both directions that founder is assigned for the whole window. If, due to recombination, two founders give rise to different sections of the window, the examinations in both directions will not agree in regions where a recombination has occurred. Where there is agreement a founder is assigned and where there is no agreement a founder is not assigned.

#### 5. Build the consensus haplotype and impute missing alleles

In the fifth step, the founder assignments defined in step 4 are used to build a consensus haplotype for the focal individual, its relevant parent and grandparent and to impute alleles for the parent and progeny. This imputation is done if an allele was inferred on the shared haplotype for two of the three individuals in a previous step.

#### 6. Multiple family phasing

AlphaFamSeq works from the oldest to the youngest individual in the pedigree. When multiple and related families are provided, the information of related individuals are accumulated in step 1 (single locus peeler) and in steps 4 and 5. The haplotypes generated for the focal individual, its parents and grandparents (steps 4 to 5) are dropped down into the pedigree if one of the focal pedigree members is related with other individuals in the pedigree.

### Examples of method implementation: Description of datasets

The algorithm was tested on a range of simulated data sets that were designed to reflect different use cases of AlphaFamSeq, and quantify the scenarios where AlphaFamSeq performs well or performs poorly. Simulated data was generated using AlphaSim [15], AlphaSeqOpt [5] and associated programs and tools. The simulation and analysis of this data involved four steps: (i) Simulation of genomes for animals in a pedigree; (ii) Simulation of whole genome sequence data; (iii) Simulation of a range of sequencing scenarios; (iv) Phasing of this sequence data with a range of algorithm parameters; and (v) Assessment of phasing accuracy. Ten replicates of each scenario were simulated and the results are the average of these replicates.

### 1. Simulation of genomes for animals in a pedigree

Sequence data were generated using the Markovian Coalescent Simulator (MaCS) [16] and AlphaSim [15] for 1,000 base haplotypes for 1 chromosome of 1M in length. The simulator used a per site mutation rate of 2.5 ×10^−8^, a per site recombination rate of 1×10^8^, and an effective population size (Ne) that varied over time in accordance with estimates for a livestock population [17]. The resulting sequences had 750,300 segregating sites in total. From the whole sequence, 700,000 (MAF=>0.001) segregating sites were selected to represent whole genome sequence data for the subsequent analysis. After the simulation of base haplotypes, AlphaSim [15] was used to drop them through a pedigree with 24 generations containing 59,194 individuals of which were 2,210 sires and 13,421 dams.

### 2. Simulation of whole genome sequence data

Sequence data were simulated for each individual by sampling sequencing reads for the 700,000 segregating sites per chromosome. This was done using a Poisson-Gamma process which allowed the number of sequence reads per locus to vary along the genome and to vary between individuals [9]. First sequenceability (γ_j_) of each of the 700,000 loci along the genome was sampled according to a gamma distribution, with shape and scale parameters equal to α and 1/α, respectively. The number of reads (*r*) per individual *i* at locus *j* was then sampled from a Poisson distribution with mean equal to *μ*_*i,j*_ = *xγ_j_*, where *x* was the targeted (average) coverage and γ_j_ was the sequenceability for the locus *j*. Each read was generated by randomly sampling one of the alleles from the two gametes of individual *i* at locus *j*, accounting for a sequencing error. All individuals in the analyses (within and across replicates) had the same shape of coverage distribution per locus (γ_j_). The α parameter in the gamma distribution was set to 4. The assumed error rate was 1‰ (ε = 0.001).

### 3. Simulation of a range of sequencing scenarios

Two sets of sequencing scenarios were simulated.

The first set of scenarios was designed to assess the performance of AlphaFamSeq for a focal individual when sequence data was available only on the focal individual, its parents and its grandparents and there was no other pedigree information. We refer to this set of scenarios as **Single Family Phasing**.

The second set of scenarios was designed to assess the performance of AlphaFamSeq when sequence data was available for several focal individuals that were interrelated in ways that are typically found in livestock populations. Interconnected pedigree information and sequence data were available for many individuals within the pedigree. We refer to this set of scenarios as **Multiple Family Phasing**.

#### Single Family Phasing

The Single Family Phasing scenarios had modest computational requirements (on average 175.95Mb of RAM and 51.1 seconds run time), which enabled the performance of the algorithm to be assessed with different values for the user-defined parameters and when the sequencing coverage of the individuals varied. To set up the Single Family Phasing scenarios, ten families, including a focal individual and their parents and grandparents, were randomly extracted from a pedigree of 59,194 individuals. The focal individuals were then phased using AlphaFamSeq using only the sequence data on individuals in the family.

We tested the impact of the allele threshold and window size on the performance of AlphaFamSeq. In these scenarios, all the 7 members were sequenced at the same coverage and we only tested 5 sequencing coverages (1*x*, 2*x*, 5*x*, 15*x*, and 30*x*). We tested six values for the allele threshold (≥ 0.60, ≥ 0.80, ≥ 0.90, ≥ 0.95, 0.99 and ≥ 0.999) and ten values for window size (1, 100, 500, 1000, 5000, 10000, 50000, 100000, 350000 and 700000). The window size represents the number of variants per each window, where “1” means that single locus phasing was performed (i.e., steps 4 and 5 of AlphaFamSeq were overpassed) and “700000” means that a single window, with all the variants in the chromosome, was used.

We also tested the performance of AlphaFamSeq when individuals were sequenced with variable coverage. In these simulations we fixed the allele probability threshold and window size to the best performing values in the previously described simulations. All combinations of six sequencing coverages were tested (0, 1, 2, 5, 15 and 30*x*). Because there were 7 members of a family (focal individual, parents and grandparents) this resulted in 6^7^ (i.e., 279,936) sequencing scenarios. We used the same 10 families as replicates and averaged their results.

#### Multiple Family Phasing

The Multiple Family Phasing scenarios were designed to test the performance of AlphaFamSeq when applied to what might be a typical livestock scenario in which multiple families are connected via a large pedigree and some members of many of these families are sequenced at different coverages. A second motivation for the Multiple Family Phasing scenarios was to test its complementarity to AlphaSeqOpt. AlphaSeqOpt is an algorithm for distributing sequencing resources across such a pedigree [5]. AlphaSeqOpt distributes sequencing resources in proportion to the genomic footprint of a focal individual and assigns some resources to the parents and grandparents of focal individuals in order to enable the sequence data of the focal individual to be phased.

To test the Multiple Family Phasing we used AlphaSeqOpt [5] to identify 500 focal individuals from the pedigree of 59,194 individuals and to distribute £317,800 worth of sequencing resources (on average 2*x*/individual) across the families of these focal individuals with each individual being sequenced at either 0*x*, 1*x*, 2*x*, 5*x*, 15*x* or 30*x*. The available budget, number of focal individuals, and number of possible sequencing coverages resulted in a total of 279,936^500^ possible ways of distributing the sequencing resources across the focal individuals and their parents and grandparents. The focal individuals were chosen by AlphaSeqOpt based on the genotype data for 2,100 markers. The assumed sequencing costs were £40 for library preparation and £80 for each 1*x* whole genome sequence. The full set of parameters for AlphaSeqOpt [5] are reported in the additional file 1. The top 500 focal individuals, their parents and grandparents encompassed a pedigree of 1,589 individuals that were sequenced at a range of coverages. This data was phased using AlphaFamSeq with the allele threshold set to equal or greater than 0.90 and the window size to 100000 variants. These parameters were chosen based on the results of the earlier scenarios.

### 4. Assessment of phasing accuracy

Phasing was performed for each scenario using the parameters previously described. The performance of AlphaFamSeq was measured as: (i) the percentage of alleles across all variants that were correctly phased (**%Correct**); and (ii) the percentage of alleles across all variants that were incorrectly phased (**%Error**). Most of the results were presented as %Correct and %Error for the focal individual. Where specified in the results section, these statistics were measured also for the parents and grandparents. Moreover the %Correct and %Error were calculated for the imputed genotypes to test the effect of the user-parameters in imputation of the homozygotes and heterozygote genotypes. In the multiple family imputation, correlations between true and imputed genotypes were calculated and binned by minor allele frequency.

## Results

AlphaFamSeq correctly phased high percentages of alleles (%Correct) and incorrectly phased very low percentages (%Error). AlphaFamSeq always performed well when sequencing coverage was high and our aim was to identify the conditions under which it performs well at low or intermediate coverage.

The results show four things: (i) in Single Family Phasing, allele threshold had a big effect and window size had a very small and non-monotonic effect; (ii) when all individuals in a single family were sequenced at different coverages the best possible %Correct reached 90% of the possible maximum (98.9%) at ~1/6 of the maximum aggregate coverage; (iii) adding sequence information from other related individuals increased the %Correct; and (iv) imputation performance was good across all allele frequencies (average correlation by marker of 0.94), except for a slight decrease at for very low frequencies (≤0.01 MAF).

### Impact of AlphaFamSeq parameters on performance of Single Family Phasing when all individuals were sequenced at the same coverage

In Single Family Phasing, allele threshold had a large effect and window size had a small effect. The %Correct ranged from 0.47% to 99.06% and the %Error ranged from 0.0001% to 8.68%.

#### User-defined allele probability threshold

The allele threshold affected both the %Correct (Figure 2) and %Error (Figure 3) more when individuals were sequenced at low coverage. Relaxing the allele thresholds (i.e., moving leftwards on the x-axis of Figure 2 and Figure 3) increased both the %Correct and the %Error for the focal individual.

**Figure 2.**
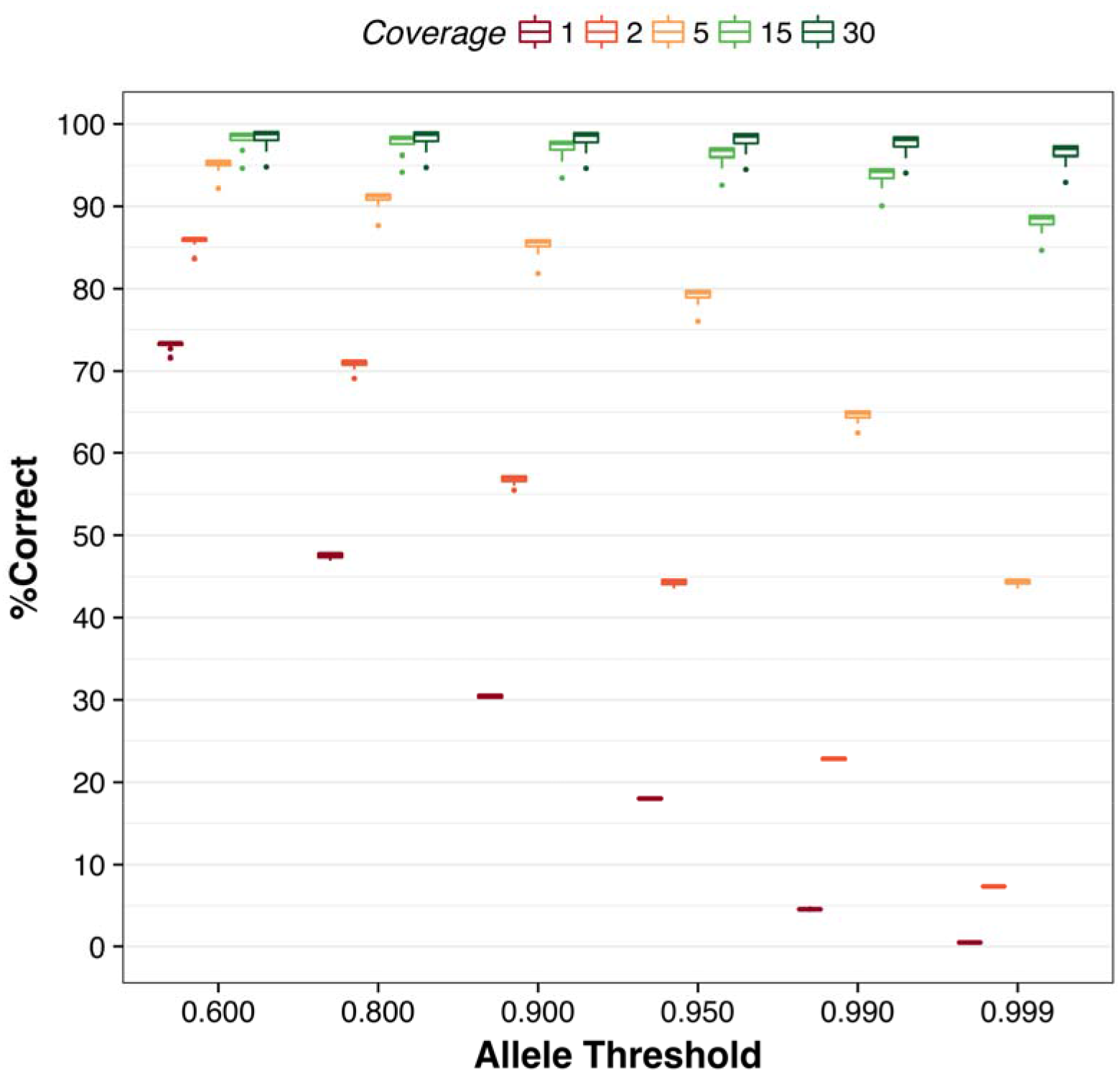
Average effect of different allele thresholds and sequencing coverage on the percentage of correctly phased alleles (%Correct) for the focal individuals when all the seven family members are sequenced at the same coverage (1x, 2x, 5x, 10x, 15x, or 30x) within each analysis.

**Figure 3.**
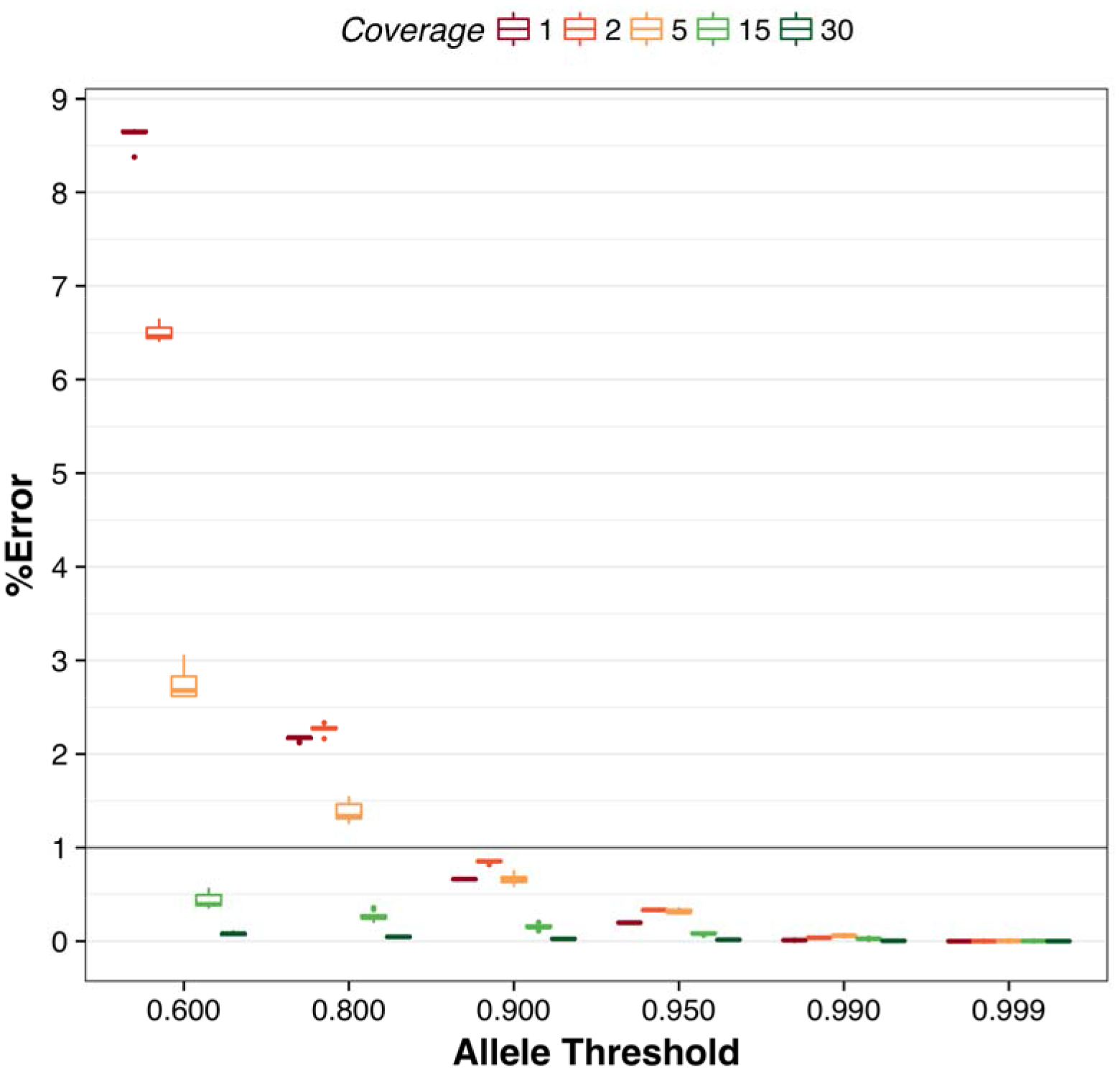
Average effect of different allele thresholds and sequencing coverage on the percentage of incorrectly phased alleles (%Error) for the focal individuals when all the seven family members are sequenced at the same coverage (1x, 2x, 5x, 10x, 15x, or 30x) within each analysis.

Unsurprisingly, the impact of relaxing the allele threshold depended on the sequencing coverage. Relaxing the allele threshold had higher impact when the individuals in the family were sequenced at low-coverage (1*x*) than when they were sequenced at high-coverage (30*x*). Relaxing the highest (0.999) to the lowest allele threshold (0.6) increased the %Correct in the focal individual by 72.58% for a family sequenced at 1x and by 1.81% for a family sequenced at 30x (Figure 2), and increased the %Error by 8.63% for a family sequenced at 1x and by 0.08% for a family sequenced at 30x (Figure 3).

Figure 2 and Figure 3 show that an allele threshold of 0.90 gave high %Correct and sufficiently low %Error (<1.00%). For this reason, in the next section of results we fix the allele threshold at 0.90 and explore the impact of window size for the 50 sub-scenarios of the Single Family Phasing.

#### Window size

Within a given sequencing coverage, changing window sizes produced small differences in %Correct and tiny and non-monotonic differences in %Error (Figure 4). The differences in %Correct were only visible at 5x (4.01% of differences) or higher sequencing coverage. Increasing window size increased the %Correct. This is because bigger window sizes allowed more haplotypes to be identified as being shared by the focal individual, its relevant parent and grandparent. At these coverages the differences in %Error were still tiny (≤0.18%).

**Figure 4.**
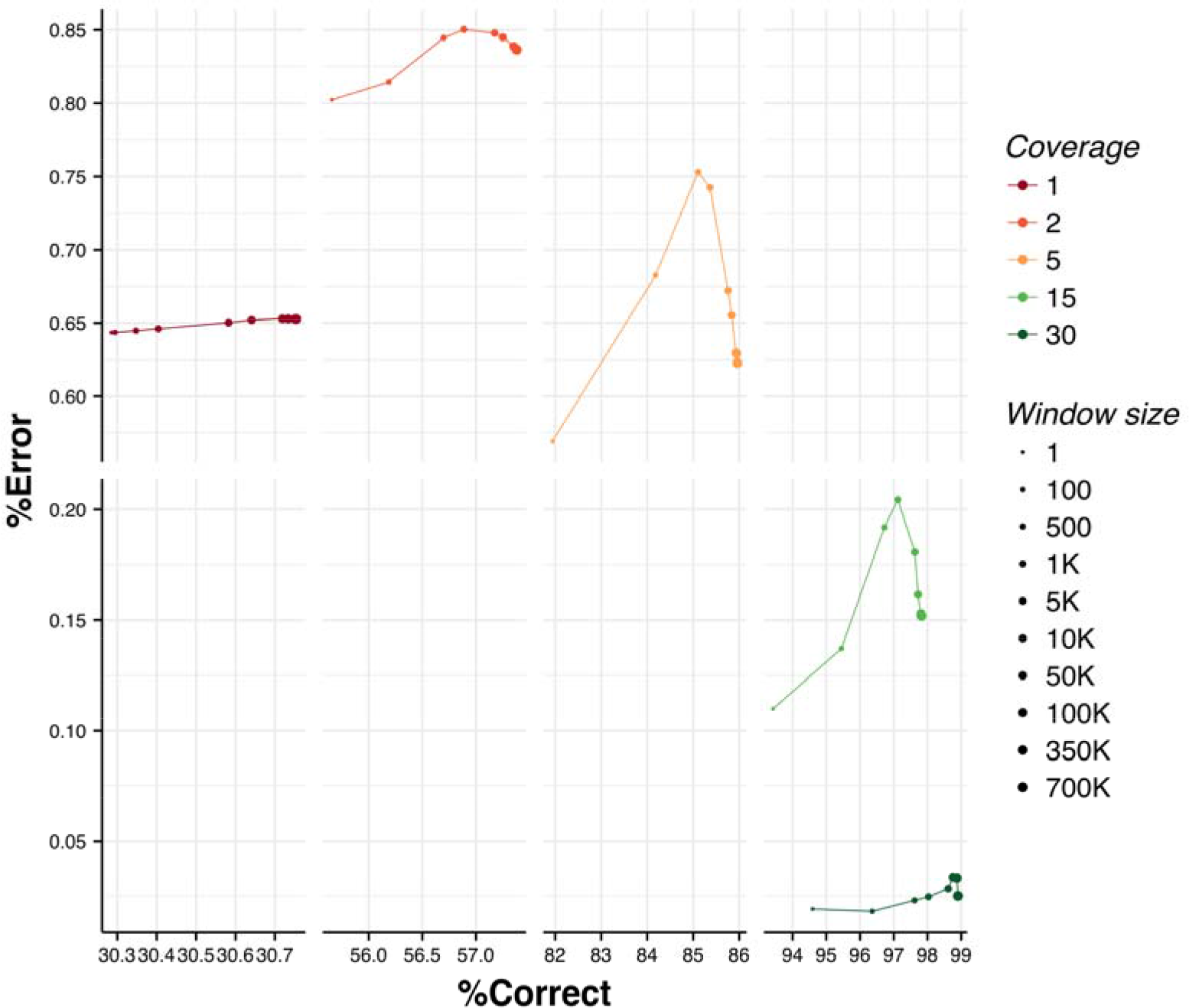
Effect of different window sizes and sequencing coverage on the percentage of correctly phased alleles (%Correct) and on the percentage of incorrectly phased alleles (%Error) for the focal individuals when all the seven family members are sequenced at the same coverage (1x, 2x, 5x, 10x, 15x, or 30x) within each analysis and the allele threshold is set to 0.90.

When a family was sequenced at 1*x*, differences were too small to see: less than 0.47% for %Correct and less than 0.01% for %Error. At 2*x* the differences were also very small: less than 1.72% for %Correct and less than 0.05 for %Error. One of the reasons for these trends is that coverage less than or equal to 2*x* on the 7 family members produced a very low number of phased alleles at heterozygote variants in a given window and thus there were not enough phased alleles to build the consensus haplotypes even if the search space (i.e., window size) was large.

Figure 4 shows that a window size of 100,000 variants gave similar %Correct and %Error to bigger window sizes but made it possible to parallelise the computation of step 4 and 5. For this reason, in the next section of results we set the window size to 100,000 variants.

#### Phasing behaviour for focal individual, parents and grandparents

AlphaFamSeq is designed to phase the sequence data of the focal individual. Some phasing is achieved for the parents and grandparents but this is a by-product of the steps taken to phase the focal individual and we expected to have less yield. Figure 5 shows that for all the sequencing coverages the focal individuals had the highest %Correct, with the parents next, and the grandparents last. The differences in %Correct between focal individual, parents and grandparents tended to decrease as the sequencing coverage increased. At 1*x* the percentage of correctly phased alleles was 30.62% (focal individual), 23.51% (parents) and 2.04% (grandparents). Whereas at 30*x* the percentage of correctly phased alleles is 98.9% (focal individual), 93.83% (parents) and 68.96% (grandparents).

**Figure 5.**
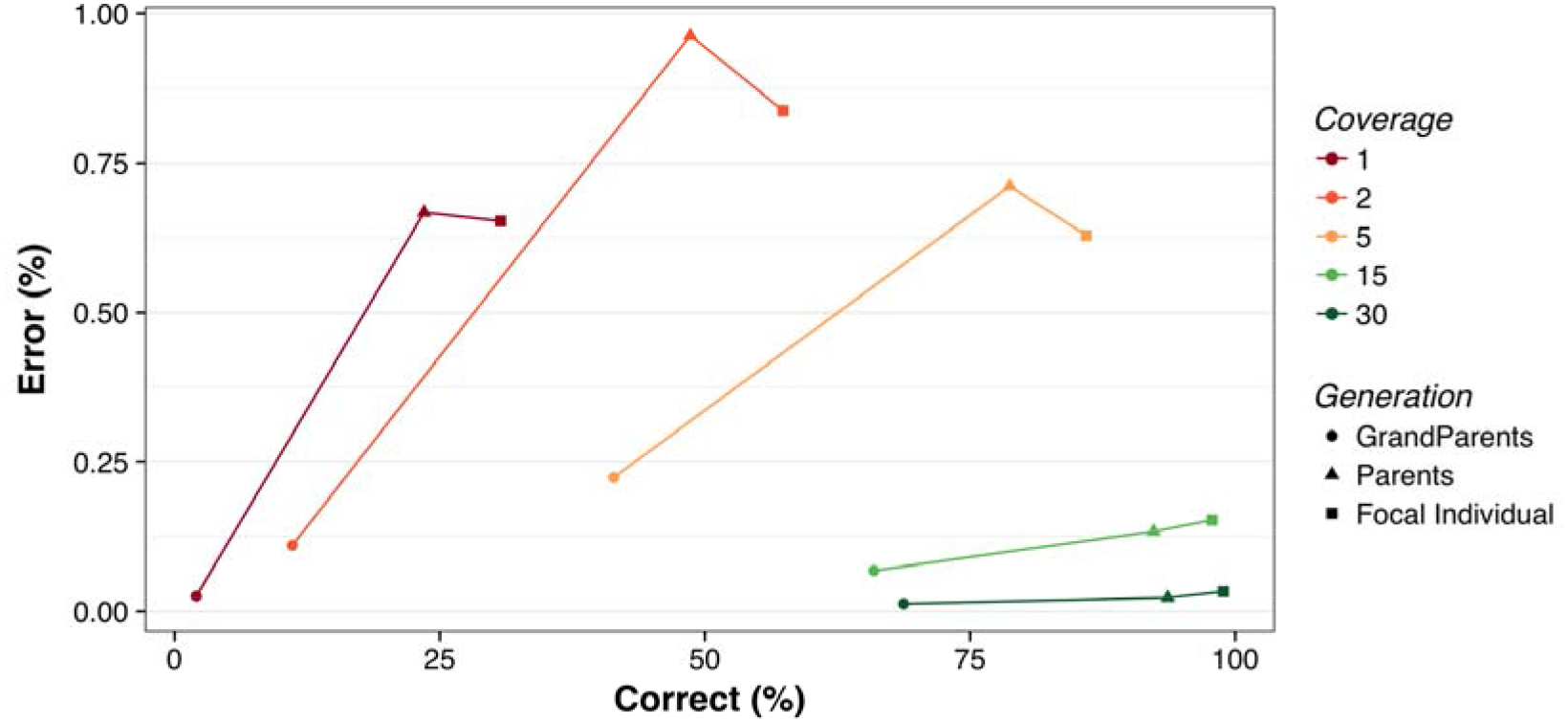
Effect of generation and sequencing coverage on the percentage of correctly phased alleles (%Correct) and on the percentage of incorrectly phased alleles (%Error) for the focal individuals when all the seven family members are sequenced at the same coverage (1x, 2x, 5x, 10x, 15x, or 30x) within each analysis, allele threshold is set to 0.90 and window size is set to 100,000 variants.

The %Error behaved differently. The grandparents always had the lowest %Error (≤0.22%), followed by the focal individuals (≤0.85%) and the parents (≤0.95%).

#### Genotype imputation

Figure 6 shows the results of imputation for the homozygote genotypes “0” and “2”, and the heterozygote genotype “1”. At all the sequencing coverages tested, the heterozygote genotypes always had the highest %Error, reaching 1.63% at 5*x* and 1.55 at 2*x*. The two homozygote genotypes had similar %Error (≤0.25%).

**Figure 6.**
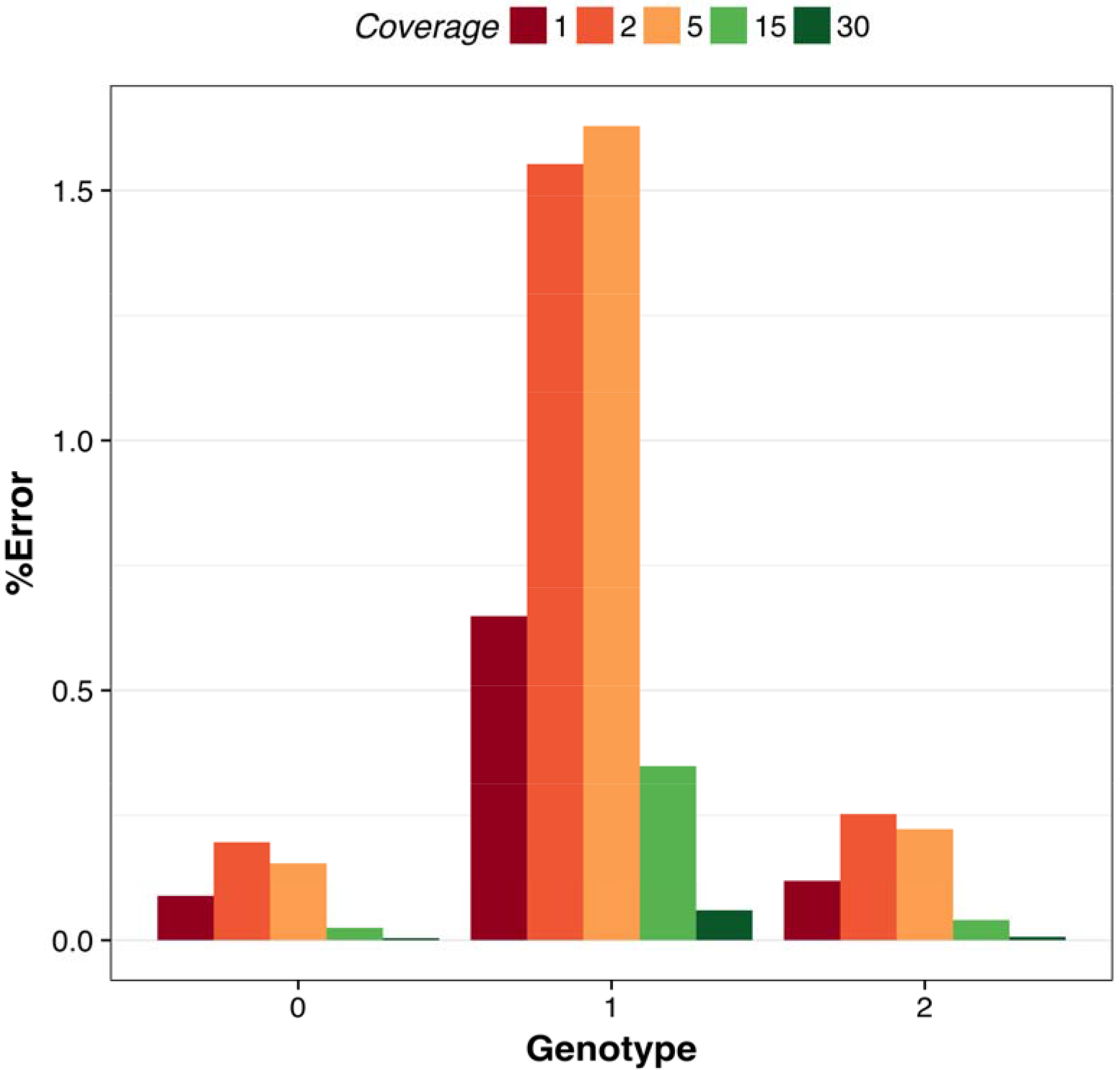
Effect of sequencing coverage on the percentage of incorrectly imputed genotypes (%Error) for the focal individuals when all the seven family members are sequenced at the same coverage (1x, 2x, 5x, 10x, 15x, or 30x) within each analysis, allele threshold is set to 0.90 and window size is set to 100,000 variants.

### Impact of AlphaFamSeq parameters on Single Family Phasing when all individuals in the family were sequenced at different coverages

When all individuals in a single family were sequenced at different coverages, the best possible %Correct reached 90% of the possible maximum at ~1/6 of the maximum aggregate coverage. Figure 7 summarizes the %Correct for each of the 279,936 scenarios of Single Family Phasing when all individuals in the family were sequenced at different coverages. The total aggregate coverage for one family of 7 members is shown on the x-axis and the %Correct is shown on the y-axis. The %Correct for the focal individual ranged from 0% to 98.9%. The %Error ranged from 0% to 3.63%. The horizontal dotted line represents the scenarios with the biggest difference in term of aggregate coverage (from 14*x* to 180*x*) that produced the same %Correct (69.6%), and highlights that the same phasing accuracy can be achieved at very different aggregate coverages. By way of example, in the scenario with 14*x* of aggregate sequencing coverage the focal individual was sequenced at 5*x*, the sire at 1*x*, the dam at 5*x*, the paternal grandsire at 0*x*, the paternal granddam at 1*x*, the maternal grandsire at 1*x*, the maternal granddam at 1*x*, while in the in the scenario with 180*x* of aggregate sequencing coverage the focal individual was not sequenced, and the parents and grandparents were sequenced at 30*x*.

**Figure 7.**
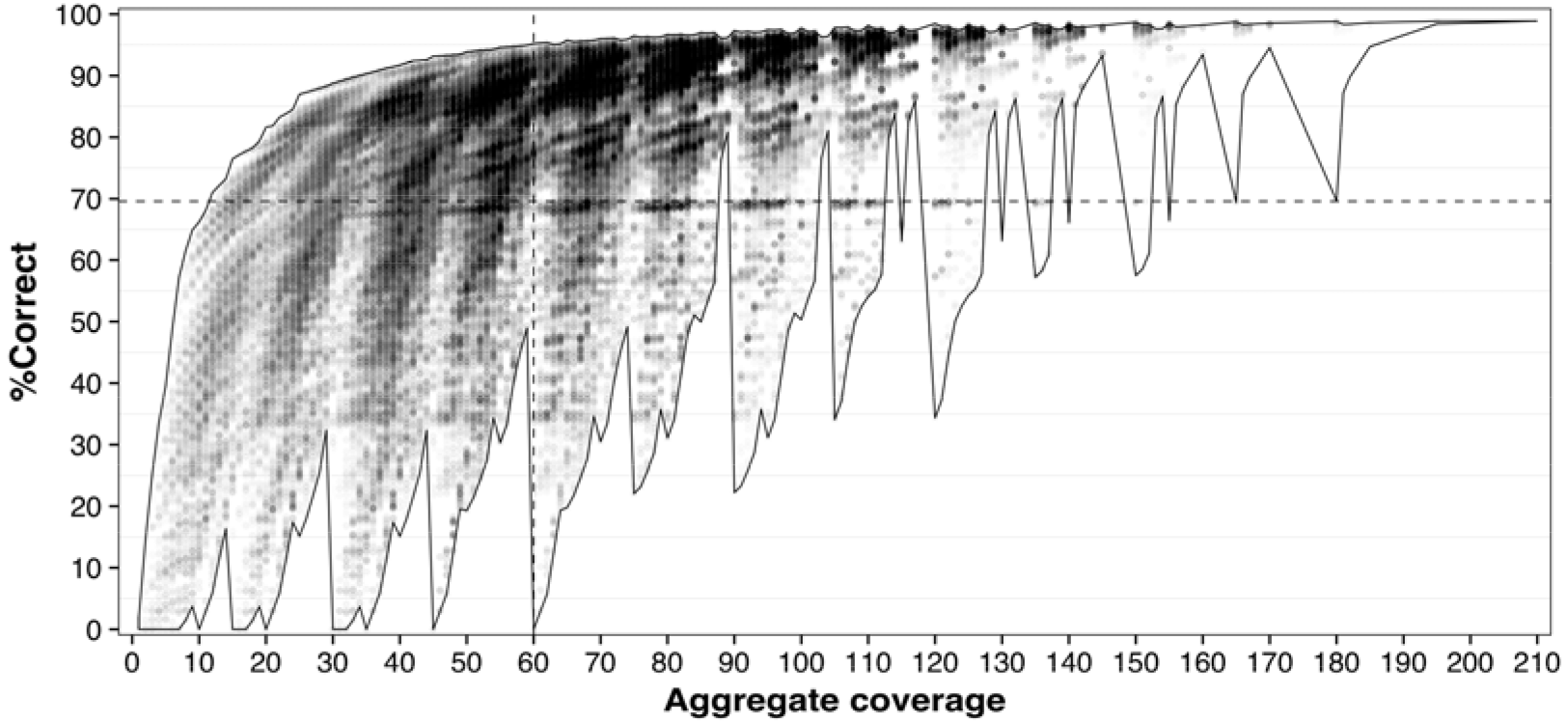
Average effect of the family aggregate coverage on the percentage of correctly phased alleles (%Correct) for the focal individuals when AlphaFamSeq is tested within independent families of 7 members. Allele threshold equal to 0.90 and window size is 100,000 variants. All the possible 279,936 combinations of sequencing depths (0, 1x, 2x, 5x, 15x and 30x) are tested.

The vertical dotted line represents the scenarios with the biggest difference in terms of %Correct (from 0% to 95.2%) at the same aggregate coverage of 60*x*, and highlights the fact that the same aggregate sequencing coverage can deliver very different phasing accuracies. Good phasing accuracy for the focal individual could be achieved with many different investments and distributions of investments. For example, by investing 100*x* aggregate coverage (the focal individual at 30*x*, the parents at 15*x*, 2 of the grandparents at 15*x*, and the other 2 grandparents at 5*x*), a %Correct of 97.75% could be achieved. In contrast sequencing all seven family members at 30*x*, more than doubled the aggregate coverage (210*x*), only produced a 1.15% increase in %Correct (98.9%).

Overall, the cloud of scenarios in Figure 7 shows that there are different combinations of sequencing coverage in a 7 member family that have different aggregate coverage, and thus costs, and give different %Correct. This cloud is taken as input by AlphaSeqOpt when it is optimising the distribution of sequencing resources across a population.

### Performance of Multiple Family Phasing

Phasing multiple families simultaneously, increased the %Correct and slightly increased the average %Error.

AlphaSeqOpt was instructed to find 500 focal individuals from the pedigree of 59,194 individuals and to assign £317,800 worth of sequencing resources to these individual and their ancestors in an optimal way. It assigned sequence resources to 1,137 individuals on 1,589 in total. Of these 176 individuals sequenced at 1*x*, 626 individuals sequenced at 2*x*, 305 individuals sequenced at 5*x* and 30 individuals sequenced at 15*x* (Figure 8A), and the total budget used was £317,720. To quantify the increase in phasing performance produced by phasing multiple connected families simultaneously, the data was phased by AlphaFamSeq using three generation pedigree that connected the multiple families and by treating each of the 500 families of the focal individuals as independent unconnected families. Based on the single family phasing results we chose an allele probability threshold of 0.90 and a window size of 100,000 variants for the analysis of the Multiple Family Phasing scenario.

**Figure 8.**
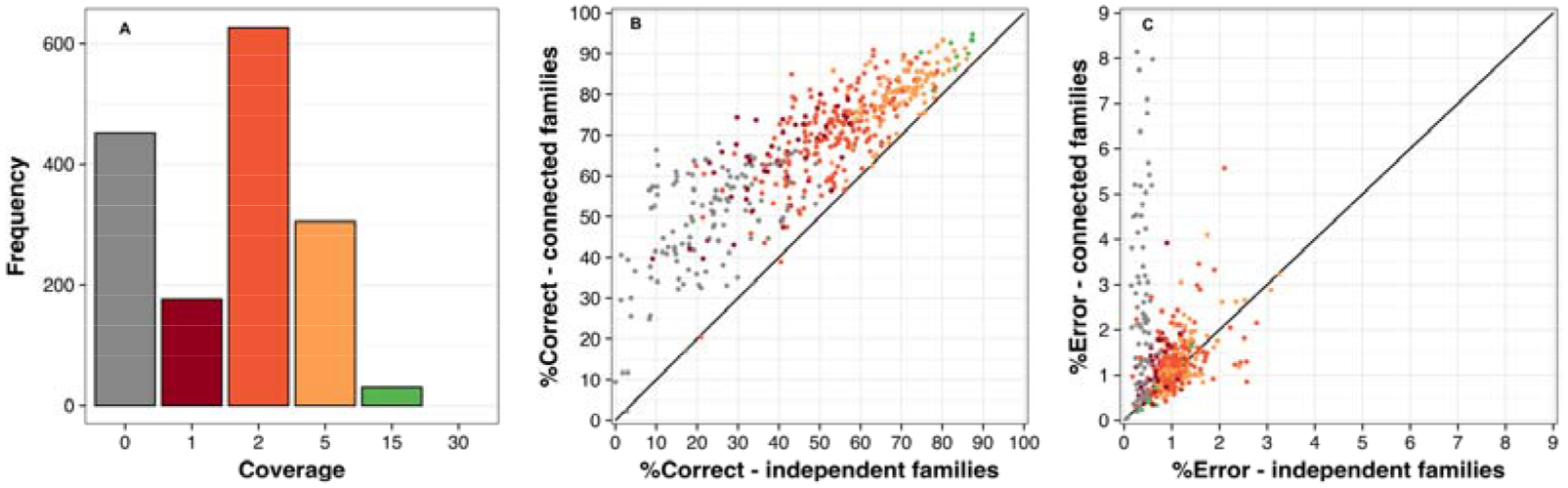
**A)** Distribution of sequencing coverage for 1,589 individuals in the pedigree when a budget of £317,720 was invested to sequence the individuals. **B)** Expected correctly phased alleles (%Correct – independent families) against the realised correctly phased alleles (%Correct – connected families) for the 500 focal individuals. **C)** Expected incorrectly phased alleles (%Error – independent families) against the realised incorrectly phased alleles (%Error – connected families) for the 500 focal individuals.

The phasing results for the focal individuals are reported in Figure 8B and Figure 8C. Figure 8B shows %Correct for the focal individuals when treating the families as independent and unconnected is shown on the x-axis. The %Correct for the focal individuals when treating the families as connected is shown on the y-axis. The average %Correct for the focal individuals increased from 47.08% to 65.25% when the families were treated as connected. 486 of the 500 focal individuals had a higher %Correct and the greatest increase in %Correct was from 10.03% to 66.34%. The focal individuals who gained most from connecting families were the 133 focal individuals not sequenced (from 22.70% to 48.15%), followed by the 53 focal individuals sequenced at 1*x* (from 40.31% to 63.67%), the 206 focal individuals sequenced at 2*x* (from 51.8% to 68.33%), the 55 focal individuals sequenced at 5*x* (from 69.67% to 79.93%) and finally the 9 focal individuals sequenced at 15*x* (from 83.3% to 90.08%). The individual with the greatest reduction in %Correct by the connecting the families only had a small reduction (-1.58 %).

Figure 8C shows that the average %Error increased by connecting the families in the analysis from 0.84% on average to 1.51%. The greatest increase in %Error was for the 133 focal individuals not sequenced (from 0.32% to 2.39%), followed by those sequenced at 1*x* (from 0.75% to 1.14%), at 2*x* (from 1.02% to 1.21%) and at 5*x* (from 1.19% to 1.24%) whereas for the 9 focal individuals sequenced at 15*x* the average %Error decreased from 0.86% to 0.81%.

### Impact of allele frequencies on imputation results

Imputation performance was good across all allele frequencies, except for a slight decrease for very low frequencies ≤0.01 - Figure 9). On average the correlation by marker between true and imputed genotype was 0.94. As we would expect, the accuracy of imputation is low at low allele frequencies: 0.63 at MAF bin ≤0.01.

**Figure 9.**
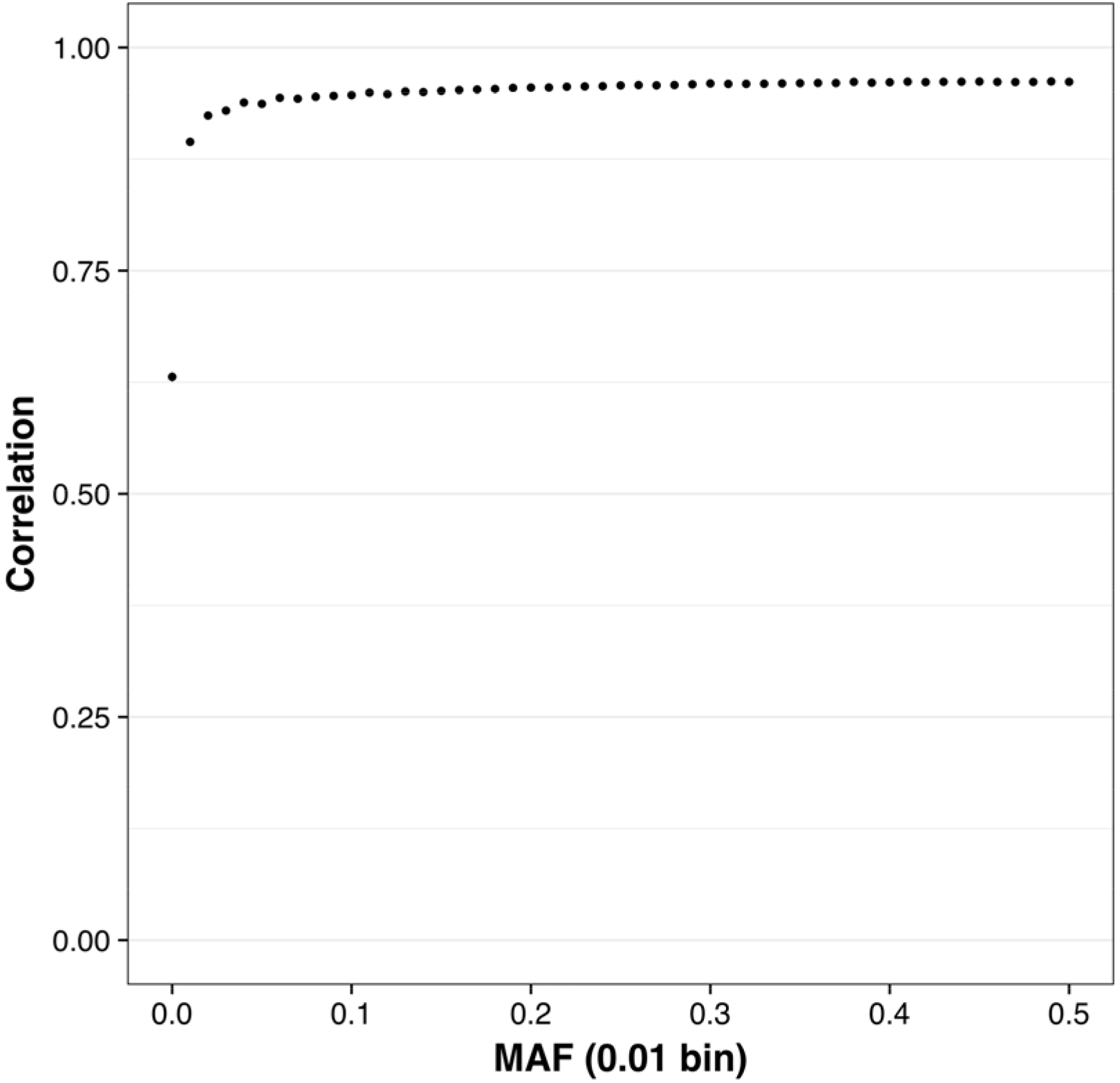
Performance of AlphaFamSeq on correlation by markers binned by minor allele frequency (MAF).

## Discussion

In this paper we developed a family-based phasing algorithm for variable-coverage sequence data that first minimises phasing errors and then maximises the proportion of alleles phased. This algorithm performed well when variable coverage sequence data was present for an individual and their parents and grandparents. Accuracy was increased when additional information was available on more distant relatives. Performance was stable across a range of user-defined parameters. We next discuss the algorithm’s performance, possible improvements to it, and its place in a toolkit for imputing and phasing livestock sequence data.

### The performance of the algorithm

In the single family phasing scenarios where focal individuals were phased based on only their sequence data and that of their parents and grandparents, we found that increasing the coverage increased our accuracy, measured by the percentage of correctly phased alleles (%Correct) regardless of the parameters. At 30*x* the average %Correct was 97.6% and the average %Error was 0.03%.

Scenarios with extremely high or extremely low allele thresholds performed poorly in families sequenced at low coverage. For example, in single families sequenced at 1*x*, a threshold that was too strict (i.e. 0.999) resulted in almost no phased alleles (0.47 %Correct), whereas a threshold that was too relaxed (i.e. 0.6) showed a high %Error (8.68%). For this reason we choose to use an allele threshold of 0.9, because it was the one that give a sufficiently low %Error (<1%) and a sufficiently high %Correct (>30%). The allele threshold had a lower impact on scenarios with family sequenced at high coverage. The reason is that individuals sequenced at high coverage have enough observed reads for the variants and so it is much easier to accurately phase their alleles and to impute any missing alleles based on information from family members. On the other hand, individuals sequenced at low coverage do not have enough observed reads to accurately phase the alleles. Decreasing the allele threshold to low values increases the chance of phasing the alleles, but increases the chance of phasing them incorrectly.

The size of the imputation window had a very small effect in a single family. When a family was sequenced at ≤2*x* differences were too small to see. When a family was sequenced at 5*x* coverage or higher increasing the window size increased the %Correct. One of the reasons for this trends is that coverage less than or equal to 2*x* on the 7 family members produced a very low number of phased alleles at heterozygous loci in a given window and thus there were not enough phased alleles to build the consensus haplotypes even if the search space (i.e., window size) was large. The highest %Error occurred at intermediate window sizes. One of the possible explanations is that, at each iteration, AlphaFamSeq was set to relax the allele threshold to a greater degree, from a value of 1.00 in the first iteration to a value of 0.90 in the final iterations. Thus, in the early iterations the algorithm used highly informative variants and, with large window sizes, there were sufficient informative variants to build and to impute highly accurate haplotypes for the focal individual and its parents and grandparents. With small window sizes relaxing the allele threshold increased the number of informative genotypes for each window and thus enabled consensus haplotypes to be built and imputed, however the accuracy of these haplotypes was lower, which increased the %Error.

Results on single families sequenced at fixed coverage lead us to believe that it is better to sequence individuals at high coverage to increase the phasing results. But results in Figure 7 show that our algorithm performs well with variable coverage. In fact, the clouds of results in Figure 7 shows that the maximum %Correct was 98.9% when all the 7 family members were sequenced at 30*x* (aggregate of 210*x*), but that a similar accuracy could be obtained with only an aggregate of 122x across a family, which is on average 17.4*x* per individual.

Our algorithm also performs well when multiple families are phased in the context of a larger, connected pedigree. Connecting families sequenced at variable coverage increased the %Correct for the focal individuals. In the scenario with an average sequencing coverage of 2*x* the %Error were still acceptable (on average 1.52%), although it increased substantially for some individuals without sequence data (up to 11.3 %Error). Moreover, imputation was good at all allele frequencies (average correlation by marker of 0.94), although there was a slight decrease for low allele frequency (≤0.01 MAF).

### Possible improvements to the algorithm

Sequence data is noisy for a variety of reasons (i.e., quality of the reference genome [18,19], misalignment of the reads [20], index switching [21,22], reference allele bias [23]). Low coverage sequence data also comes with the risk of calling genotypes incorrectly due to a small number of reads (i.e., heterozygous genotypes that are called as homozygous because only one of the alleles has read information). Being able to call alleles at a high accuracy while minimizing the number of incorrectly called alleles is essential for using low coverage sequence data. In populations that share segments identical-by-descent, there is an advantage in performing the allele- and genotype-calling accumulating the sequence information of related individuals, as AlphaFamSeq does. This increases the accuracy of phasing for individuals that are sequenced at low-coverage.

In the first step of AlphaFamSeq we utilize an iterative peeling algorithm to initially pass and accumulate read information between relatives [13]. This peeling algorithm works only on a single allele at a time, utilizing family relationships, and does not use linkage information. We anticipate that it would be possible to increase the yield of the initial iterative peeling steps by using a multi-locus version of the iterative peeling algorithm, such as LDMIP [24]. Multi locus peeling has the advantage of taking linkage information into account, allowing for a more accurate sharing of read information. However, currently LDMIP does not scale well to whole chromosome sequence data, and so an approach such as that was not used. Moreover, the computational cost of peeling algorithms increase when the complexity and size of the pedigree increases.

The sequence data use by AlphaFamSeq are the observed number of reads for the reference and the alternative alleles. One of the possible improvements in the algorithm is to use the information from the raw sequence reads (i.e., those stored in bam or sam files [25]) as input data. These raw sequence reads have the advantage that they may cover two or more heterozygote variants and so provide information on the physical linkage of these variants. This linkage information can be used to improve the phasing accuracy and reduce the switching errors [26–28]. An extension of this would be to use some long read information from the single-molecule long-read sequencing [28]. This technology can generate reads which are much longer compare to the reads length of the NGS, but on the other hand, the sequencing errors are high [29]. Modifying AlphaFameSeq to handle such errors and utilise long read information would increase its phasing accuracy.

In our simulations we tested only small pedigrees of related individuals and we only supplied simulated sequence data to AlphaFamSeq. The incorporation of more sequence data on more animals and incorporation of cheaper genotype data on many animals would improve the phasing results. Genotype data are available nowadays for large numbers of animals, and even if they represent a small portion of SNP of the whole genome sequence, they can help to reconstruct the haplotypes of not sequenced individuals or individuals sequenced at low coverage.

### An overall strategy for imputing sequence data for whole populations and the role of AlphaFamSeq

We believe that the key to perform whole genome sequencing for large livestock populations can be broken up into four steps. First, animals that carry a large proportion of haplotypes in the population need to be identified and sequenced, potentially at variable coverage to optimise use of sequencing resources. Second, the sequence data needs to be phased to provide a reference panel for downstream imputation. Third, these two steps can be complemented with the judicious use of low-coverage sequence on some individuals. Fourthly, an imputation algorithm needs to be used to pass the phased haplotypes to the remaining individuals in the population. This sequencing strategy stems from the fact that cheap genotype arrays are a fraction of the cost of sequencing individuals, and livestock populations have a high degree of relatedness allowing for large shared haplotypes and enabling high accuracy imputation.

AlphaSeqOpt [5] solves the first problem by determining which individuals in a population should be sequenced and the coverages that they should be sequenced at. AlphaFamSeq solves the second problem by providing a set of reference haplotypes that can be used for downstream imputation. The last problem could be handled by either heuristic (e.g. AlphaImpute [6], Findhap [7], Fimpute [8]) or probabilistic (e.g. MaCH [9], Beagle [10,11], Impute2 [12]) imputation algorithms or a combination of both [30]. We discuss these steps in more detail below.

The first step of sequencing is to determine which animals (i.e., focal individuals) contain a large number of the haplotypes shared with many other animals in the population and determining the optimal distribution of sequencing resources across these focal individuals and their ancestors (parents and grandparents) according to the genomic footprint of the focal individuals on the population. Methods such as AlphaSeqOpt [5] use phased genotype data from genotype arrays to identify high-frequency haplotypes in the population and identify focal individuals who carry these haplotypes. Part of the sequencing resources is then spent on the focal individual and their parents and grandparents to genotype and phase the haplotypes of the focal individuals. To determine how to spend sequencing resources AlphaSeqOpt requires a function that estimates the phasing accuracy for a focal individual depending on the individual’s coverage and that of their family. AlphaFamSeq fills this gap by providing a way to estimate the expected phasing accuracy for individuals based on a family’s coverage.

The next step of our sequencing strategy is to take the variable coverage sequence information and phase the haplotypes of focal individuals. AlphaFamSeq, as described in this paper, performs this step and can phase alleles for focal animals. The high accuracy haplotypes generated by AlphaFamSeq are key for downstream imputation.

The two steps described above could be complemented by a sequencing strategy that we call LCSeq. LCSeq aims to generate accurate sequence for the haplotypes in the population. LCSeq sequences individuals at low-coverage and assembles high-coverage sequence information for every haplotype by accumulating the low-coverage sequence data from the genome segments that are shared between many individuals to derive the ‘consensus haplotypes’. LCSeq uses the consensus haplotypes to impute the sequence data of the individuals. Accurate derivation of the consensus haplotypes is critical for phasing and imputation accuracy. Efficient derivation of the consensus haplotypes requires distribution of sequence data across the population such that a maximal number of haplotypes are sequenced to a coverage (e.g., 20x) that is sufficient to enable them to be phased accurately, which requires an algorithm such as that developed by Ros et al. [31]. Within the context of LCSeq, AlphaFamSeq can provide set of accurate starting haplotypes from which many of the alternate haplotypes carried by individuals sequenced at low coverage can be derived. AlphaFamSeq will increase the numbers of haplotypes accurately phased by consensus because any individual that is sequenced at low coverage and who shares haplotypes with a focal individual will have those haplotypes phased and many of the other haplotypes that it carries will be phased either by being the complement of a shared haplotype or by low-coverage sequence reads.

The fourth step of our sequencing strategy involves taking the phased reference haplotypes generated by AlphaFamSeq, or LCSeq, and imputing them to the remaining individuals in the population. Several methods or combinations of methods could be used for this (e.g. AlphaImpute [6], Findhap [7], Fimpute [8], MaCH [9], Beagle [10,11], Impute2 [12]). The availability of a pre-phased set of individuals or haplotypes greatly decreases the computational time of these algorithms and increases their accuracy [11,12,30]. AlphaFamSeq, with its low percentage of alleles that incorrectly phased and high percentage of alleles that are correctly phased can provide such pre-phased data.

## Conclusions

This paper describes a family-based phasing algorithm, for variable-coverage sequence data, that first minimises phasing errors and then maximises the proportion of alleles correctly phased. We tested the algorithm in several scenarios that are typical of those found in livestock breeding and genetics. It can work within a given family separately, can work with multiple families simultaneously by utilising pedigree information that connects them and can handle variable coverage sequencing data. It gives low phasing error rates and, if there is sufficient sequence information and haplotype sharing amongst individuals, it can give a high yield of phased alleles. We envisage that AlphaFamSeq will be one of a number of essential tools that would underpin an overall strategy for generating highly accurate sequence data on whole populations at low cost. The role of AlphaFamSeq in this overall strategy is to provide very accurately phased haplotypes on focal individuals, who are individuals whose haplotypes are very common in the population.

## Supporting data

Programs to simulate the true genotypes (AlphaSim version 1.08), the sequence reads (SimulateSequenceReads), to optimally distribute the sequencing resources (AlphaSeqOpt version 1.00) and perform phasing (AlphaFamSeq version 1.00) used in this paper are available from the AlphaGenes website [32].

## Authors’ contributions

JMH conceived the algorithm and supervised the study. MB further developed the algorithm, implemented it in software, performed all of the analysis and wrote the first draft of the paper. RRF, SG, GG and AW contributed to the development of the method, the interpretation of the results and to the writing of the paper.

## Acknowledgements

The authors acknowledge the financial support from the BBSRC ISPG to The Roslin Institute “BB/J004235/1”, from Genus PLC and from grant numbers “BB/M009254/1”, “BB/L020726/1”, “BB/N004736/1”, “BB/N004728/1”, “BB/L020467/1”, “BB/N006178/1” and Medical Research Council (MRC) grant number “MR/M000370/1”. This work has made use of the rosources provided by the Edinburgh Compute and Data Facility (ECDF) (http://www.ecdf.ed.ac.uk). The authors thank Dr Andrew Derrington (Scotland, UK) for assistance in refining the manuscript.

John Hickey would like to acknowledge the outstanding contribution by Robin Thompson to the fields of statistics, animal breeding and plant breeding. Robin has been a great teacher, mentor, friend and inspiration to him over many years.

## Additional file

File name: Additional file 1 Format:.txt

Title: Parameters used to run AlphaSeqOpt.

Description: parameters used to run AlphaSeqOpt (version 1.00). The parameters are fully described in www.alphagenes.roslin.ed.ac.uk/alphasuite-softwares/alphaseqopt/

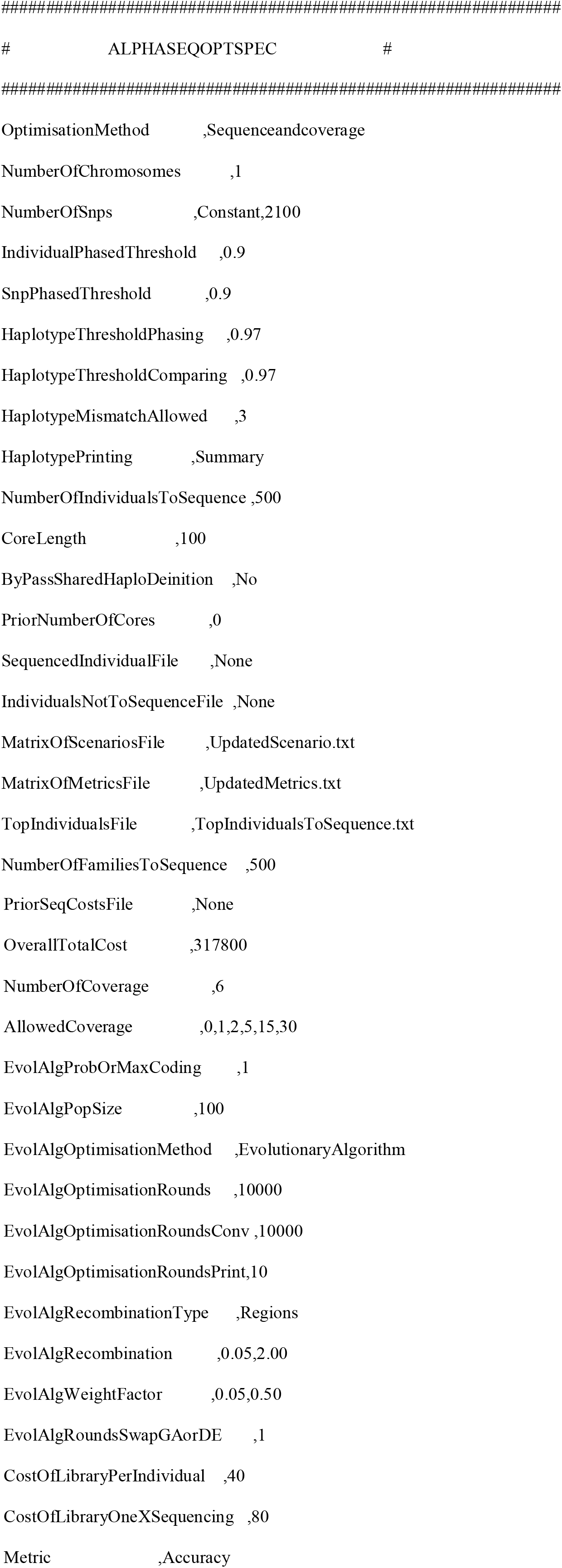

